# Multiple species drive flexible lake food webs with warming

**DOI:** 10.1101/499400

**Authors:** Timothy J. Bartley, Tyler D. Tunney, Nigel P. Lester, Brian J. Shuter, Robert H. Hanner, Kevin S. McCann

## Abstract

1. Climate change is rewiring the food webs that determine the fate of diverse ecosystems. Mobile generalist consumers are responding to climate change by rapidly shifting their behaviour and foraging, driving food webs to flex. Although these responsive generalists form a key stabilizing module in food web structure, the extent to which they are present throughout whole food webs is largely unknown.
2. Our goal was to examine if the thermal preferences of multiple species drive predictable behavioural and foraging responses to changes in temperature that drive flexes in food web structure.
3. We used stable isotope and catch-per-unit-effort data to investigate the habitat use and diet of four widespread and abundant species of fishes that represent different key trophic roles in food webs of boreal lakes that span a natural 7°C air temperature gradient.
4. We found significant reductions in nearshore derived carbon and nearshore habitat use with increased temperature in three of four fish species, indicating that multiple species drive flexible lake food webs with warming. We also found evidence that the response of lake trout to increased temperatures may reduce their biomass and cascade to release their preferred prey, the pelagic forage fish cisco.
5. Our results suggest that climate warming will shift lake food webs toward increased reliance on offshore habitats and resources. We argue that species across trophic levels broadly couple lake macrohabitats, suggesting that potentially stabilizing responsive consumers are present throughout food webs. However, climate change appears to limit their ability to responsively forage, critically undermining a repeated stabilizing mechanism in food webs.

## Introduction

Ongoing rapid climate change is altering species interactions and reorganizing the food webs that determine the fate of diverse ecosystems (Barton, Beckerman, & Schmitz, 2009; Kortsch, Primicerio, Fossheim, Dolgov, & Aschan, 2015; T. D. Tunney, McCann, Lester, & Shuter, 2014). As environmental conditions change, food web structure changes—or ‘flexes’—with it (McMeans et al., 2016). Identifying these changes in food web structure is central to understanding the consequences of climate change on the major energy pathways of carbon and nutrient flow in ecosystems but is challenging given that food web structure is inherently complex (Polis, 1991). Importantly, climate change is expected to ‘rewire’ food webs (Bartley et al., n.d.) by adding or removing whole trophic links (e.g., Kortsch *et al.* 2015) and by modifying the strengths of existing interactions (e.g., Barton & Schmitz 2009). These changes in the flow of energy among species are likely to have myriad consequences in ecosystems, from species to whole ecosystem function (Guzzo, Blanchfield, & Rennie, 2017; Kondoh, 2010; Křivan & Schmitz, 2003). If so, future warming may alter food webs in ways that lead to ecosystems that differ substantially from past and present conditions (Kortsch et al., 2012). Despite the potential for these drastic changes, our understanding of food webs responses to climate change and the consequences of these response remain limited.

Food webs are flexible because of the foraging responses of mobile generalist consumers that can rapidly shift their foraging behaviour in response to environmental change. Many generalist consumers move across the landscape to forage in various spatially-distinct macrohabitats (Rooney, McCann, & Moore, 2008) and can feed omnivorously across trophic levels (Thompson *et al.* 2007). As a result, these generalist consumers can respond in two key ways: by changing the degree to which they couple different macrohabitats (i.e., ‘horizontal’ shifts between energy pathways, Schindler & Scheuerell 2002; Vander Zanden & Vadeboncoeur 2002; Rooney *et al.* 2008) and by shifting their degree of omnivory (i.e., ‘vertical’ foraging shifts along an energy pathway, Barton & Schmitz 2009; Sentis *et al.* 2014; Ruiz-Cooley *et al.* 2017). Generalists that exhibit both horizontal and vertical foraging flexibility form the generalist module (*sensu* McMeans *et al.* 2016), a widely documented patterning of food web interactions (Eloranta et al., 2015; Nakano & Murakami, 2001; Polis, Anderson, Anderson, & Holt, 1997; Rooney et al., 2008) that can expand and contract food webs in response to changing environmental conditions (T. D. Tunney, McCann, Lester, & Shuter, 2012). This generalist module may be a powerful stabilizing structure, allowing mobile generalists to respond to changing densities in different habitats in a manner that buffers population dynamics (Kevin Shear McCann & Rooney, 2009; McMeans et al., 2016). Despite this potential, few studies have empirically documented the presence of fundamental module throughout food webs or how human impacts like climate change may interrupt this module.

One well-documented example of a responsive generalist consumer that drives food web flexibility is lake trout (*Salvelinus namaycush*), a common generalist top predator in lakes of boreal North America. Lake trout forage on fishes and invertebrates in both of the two thermally distinct lake macrohabitats: the nearshore (i.e., littoral) habitat and the offshore (i.e., pelagic) habitat (Vander Zanden & Vadeboncoeur, 2002). Because lake trout is cold-adapted and thermally sensitive, the foraging habits of lake trout are mediated by thermal accessibility of the relatively warm nearshore habitat (Guzzo et al., 2017; Plumb & Blanchfield, 2009). Lake trout show reduced nearshore habitat use with increasing nearshore temperature (Guzzo et al., 2017; T. D. Tunney et al., 2014). When lake trout avoid entering the physiologically taxing nearshore habitat, they forage less on nearshore resources and more on fish (T. D. Tunney et al., 2012, 2014). This reduced access to abundant nearshore prey may undermine lake trout’s ability to responsively forage to resource densities in different macrohabitats (Guzzo et al., 2017; T. D. Tunney et al., 2014). Importantly, this phenomenon may not be limited to lake trout; many fishes are generally highly mobile and flexible foragers (Dill, 1983), and boreal lakes in North America contain numerous cold-adapted fish species that may exhibit similar behavioural and foraging responses due to thermal limitations (Hasnain, Shuter, & Minns, 2013; Vander Zanden & Vadeboncoeur, 2002). In contrast, the species that prefer warmer waters may not be thermally limited and thus may not exhibit behavioural or foraging responses to warming. However, the roles that species other than lake trout play in determining how lake food webs flex are largely unexplored. To understand how whole food webs are being altered by climate change requires that we examine how many species across trophic levels and with various thermal preferences respond to warming.

Here, we use a natural climate gradient of approximately 7°C to examine how multiple species behavioural and foraging responses drive flexes in food web structure. We investigate the habitat use and diet of four widespread and abundant species of boreal shield fishes that represent different key roles and trophic levels in the food web: the cold-adapted generalist top predator lake trout, the cold-adapted pelagic planktivorous cisco (*Coregonus artedi*), the cool-adapted piscivorous top predator walleye (*Sander vitreus*), and the cool-adapted mesopredator yellow perch (*Perca flavescens*) (Coker, Portt, & Minns, 2001; Hasnain et al., 2013). We use stable-isotope-based food-web indices of nearshore feeding and trophic position along with catch-per-unit-effort based metrics of behaviour to show that several species across trophic levels and from multiple thermal guilds respond predictably to changes in temperature. We also use catch-per-unit-effort data to show that lake trout show reduced biomass index under increased temperatures, consistent with a reduction of the nearshore resource availability. Our results suggestion that multiple species comprising key trophic roles drive flexible lake food webs with warming. We argue that by studying the foraging and behavioral responses of this set of key players, we can predict how food webs will flex with climate change. We end by discussing how these changes suggest climate change will impact the function and stability of lake ecosystems, and how many generalist consumers may be a fundamental feature of food web architecture.

## Materials and Methods

### Lake Selection

The Canadian boreal shield includes hundreds of thousands of lakes that span various natural environmental gradients, including climate (Gunn, Steedman, & Ryder, 2004; Keller, 2007). We used data for 66 lakes in the province of Ontario, all of which have been used previously to study food web structure through stable isotope analysis (Dolson, McCann, Rooney, & Ridgway, 2009; T. D. Tunney et al., 2014; T. Tunney, McCann, Jarvis, Lester, & Shuter, 2018). We use lakes that have both nearshore and offshore intermediate consumers to ensure that changes in food web structure or predator behaviour are not driven by differences in the presence or absence of whole trophic groups. For 59 of our 66 lakes, we used catch-per-unit-effort (CUE) data from the Ontario Ministry of Natural Resources and Forestry (OMNRF) Broad-scale Fisheries Monitoring (BSM) Program (Sandstrom, Rawson, & Lester, 2013). In summary, for each lake, standardized fish community surveys took place one time between May and September from 2008 to 2012 using overnight sets of two types of multipaneled mesh gillnets: North American standard multipaneled gill nets (with 8 mesh sizes varying from 38 to 127 mm) and small mesh gillnets (with 5 mesh sizes from 13 to 38 mm). Netting occurred randomly throughout the lake and depending on the maximum depth of each lake, in each of up to eight depth strata: 1-3m, 3-6m, 6-12m, 12-20m, 20-35m, 35-50m, 50-75m, and >75m. Complete details can be found in Sandstrom *et al.* 2013. Catch data were not available for the seven lakes from Dolson *et al.* (2009).

### Lake Attributes

We used two measures of temperature for our analysis: average recent air temperature in °C and growing degree days > 5 °C (Table S1). Our primary temperature variable was average recent air temperature, which was calculated as the average hourly air temperature in °C for the 30 days prior to field sampling because it corresponds to the time period reflected by the isotopic signature of fish muscle tissue (Peterson & Fry, 1987). This temperature variable was derived from FetchClimate (Grechka et al., 2016) using the latitude and longitude of each lake. In addition to the average recent air temperature, we corroborated our results with and growing degree days above 5°C from 1981 to 2010 (provided by the Ontario Ministry of Natural Resources and Forestry), which measures the accumulation of heat over time. We also account for several physical and chemical factors that have been previously identified as important drivers of habitat coupling or trophic position in boreal shield lakes (Table S1, all provided by the Ontario Ministry of Natural Resources, Dolson *et al.* 2009; Tunney *et al.* 2014, 2018): lake surface area in hectares, mean lake depth in metres, Secchi depth in metres, and total phosphorus in μg • L^-1^, and lake shape (calculated the shoreline development index): 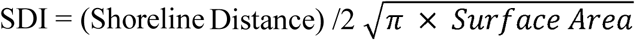 (equation 1).

### Species Selection

We used four species that are typical of their respective trophic groups. Stable isotope data for these species was available for 40 or more lakes and the trophic level, thermal classifications, and habitat preferences for these species are known (Coker et al., 2001; Hasnain et al., 2013). Lake trout (*Salvelinus namaycush*) is the most common top predator of offshore habitats in boreal shield lakes, and much research on boreal shield lake food webs has focused on this coldwater species (Dolson et al., 2009; Guzzo et al., 2017; Plumb & Blanchfield, 2009; T. D. Tunney et al., 2014). Cisco (or lake herring, *Coregonus artedi*) is one of the most common cold-adapted planktivores and is a common prey item for lake trout and walleye. Walleye (*Sander vitreus*) is a common cool-adapted piscivore and popular sport fish that is present in lakes across the boreal shield. Yellow perch (*Perca flavescens*) is a widespread and abundant cool-adapted mesopredator that is consumed by a wide variety of predatory fishes throughout its range, including both lake trout and walleye. These four species comprise a large portion of the average catch in the 59 lakes that we use here from the OMNRF’s BsM surveys (see Supplementary Information). The number of individual for each species sampled for stable isotope analysis in each lake varied from two to 21 (mean 13.5) for lake trout, three to 32 (mean 17.0) for walleye, one to 20 (mean 10.8) for cisco, and one to 18 (mean 9.1) for yellow perch. For both lake trout and walleye, only individuals greater than 250 mm were used for stable isotope analysis because these species are known to show ontogenic diet shifts (Galarowicz, Adams, & Wahl, 2006; Mittelbach & Persson, 1998; Sherwood, Pazzia, Moeser, Hontela, & Rasmussen, 2002).

### Food Web Metric Calculations using Stable Isotopes

As in many previous studies, we used stable isotopic signatures from our four fish species and baseline invertebrates to calculate both the nearshore carbon index (based on the proportion nearshore carbon, e.g., Tunney *et al.* 2014) and trophic position (David M Post, 2002; Vander Zanden, Casselman, & Rasmussen, 1999; Vander Zanden, Shuter, Lester, & Rasmussen, 1999). Collection and processing methods for stable isotope data can be found in Dolson *et al.* 2009; Tunney *et al.* 2014, 2018). We used baseline invertebrates from the nearshore and offshore zones to account for variability in isotopic signatures across lakes. We incorporated data for multiple trophic groups into both our nearshore and offshore baseline isotopic signatures to reduce the number of estimated baseline values required for our analysis and to increase the sample size for our baseline isotopic signatures. We corrected all δ^13^C signatures using C:N ratios as δ^*13*^C_corr_ = δ^*13*^C_raw_ + (−3.32 + 0.99 * CN) (equation 2), where δ^*13*^C_corr_ is the corrected δ^13^C signature, δ^*13*^C_raw_ is the raw δ^13^C signature, and CN is the C:N ratio of that tissue sample (David M. Post et al., 2007). For lakes that were missing either nearshore or offshore baseline isotopic signatures, we estimated the δ^13^C and δ^15^N signatures of the missing baseline using the available baseline and simple linear regression between the baselines across lakes (see Supplementary Information).

We used two source mixing models to estimate the nearshore carbon index and the trophic position of each species in each lake based on their relative isotopic signatures (David M Post, 2002). We calculated the nearshore carbon index for each fish species as NCI_fish_ = (δ^13^C _fish -_ δ^13^ C_osb_)/ (δ^13^C_nsb -_ δ^13^ C_osb_) (equation 3), where NCI_fish_ is the nearshore carbon index in the diet of a fish species, δ^13^C_fish_ is the average δ^13^C signature for that fish species, δ^*13*^C_osb_ is the average or estimated δ^13^C signature for all offshore baselines (i.e., mussels and/or zooplankton), and δ^13^C_nsb_ is the average or estimated δ^13^C signature for all nearshore baselines (i.e., snails and/or aquatic insect larvae). The nearshore carbon index is similar to the proportion nearshore carbon used by others (e.g., Tunney *et al.* 2014) but is not constrained between zero and one. This approach minimizes data transformation requirements of the stable isotopic signatures and increases the number of lakes in our analyses while still allowing us to qualitatively understand changes in the relative contribution of nearshore resources to fish diets.

Based on the nearshore feeding index, we estimated the trophic position of each species as TP_fish_ = 2 + (δ^15^ N _fish_ - (δ^15^N _nsb_ x NCI _fish_ + δ^15^ N _osb_ x (1 - NCI _fish_)))/3.4 (equation 4), where TP_fish_ is the trophic position of a fish species, NFI_fish_ is the nearshore carbon index for that fish species, δ^15^N_fish_ is the average δ^15^N signature for that fish species, δ^15^N_nsb_ is the average or estimated δ^15^N signature for all nearshore baselines (i.e., snails and aquatic insect larvae), δ^15^N_osb_ is the average or estimated δ^15^N signature for all offshore baselines (i.e., mussels and zooplankton), 3.4 is the assumed increase in δ^15^N due to fractionation (David M Post, 2002), for each trophic level, and 2 is the assumed trophic position of the baseline invertebrates.

### Behaviour Metrics and Biomass Index Using Catch-per-unit-effort

We used catch-per-unit-effort data for each depth stratum (Sandstrom *et al.* 2013) to calculate a weighted average mean depth of capture for each species as 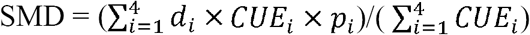 (equation 5), where SMD is the mean depth of capture of a fish species, CUE_*i*_ is the CUE of that species for depth stratum *i, p*_*i*_ is the proportion of the lake in depth stratum *i*, and *d*_*i*_ is the depth of at the center of the depth range for stratum *i* (2m for stratum 1, 4.5m for stratum 2, 9m for stratum 3, and 16m for stratum 4). We only used depth strata 1 through 4 to calculate mean depth of capture because these are the only depth strata that are sampled with both small-mesh and large-mesh gear (Sandstrom et al., 2013). The CUE data that we use here is the number of fish per 100 metre nights. For CUE biomass index, we used the kg of fish per gang per 100 metre nights, calculated by summing the area-weighted CUE (number of fish per 100 metre nights) for each depth stratum and then multiplying by the mean weight of that fish species.

### Statistical Analyses

To test for the effect of temperature on each response variable (nearshore carbon index, trophic position, mean depth of capture, nearshore/offshore presence, and the log-transformed CUE biomass index of each species), we used a two-step approach. Firstly, to account for morphometric or chemical factors known or suspected to influence the foraging and behaviour of these fish species, we first constructed multiple regression models for each response variable and lake size in hectares, lake shape (SDI), Secchi depth in m, mean lake depth in m, and total phosphorous in μg•L^-1^. We then compared all subsets of each full multivariate regression model using AIC. We selected the best model as the model with the smallest number of explanatory variables from the set of models within 2 AIC points of the lowest AIC value (Aho *et al.* 2014, see Supplementary Information for best models). Secondly, to test the effect of each temperature variable, we took the best model for each response variable and ran a multiple regression that included the temperature variable (Figs 2, 3, and 4, Table S3). To meet statistical requirements, we used log_10_ transformed lake surface area and natural log transformed mean lake depth and total phosphorous. We removed all data points with a Cook’s distance greater than 1 from their respective regression models, and we used variance inflation factor to check for inflated coefficient estimates in multiple linear regressions. All regression analyses were performed in the R statistical language (v3.2.3), and model selection was conducted using the R package “glmulti” (Calcagno, 2013).

**Figure 1.**
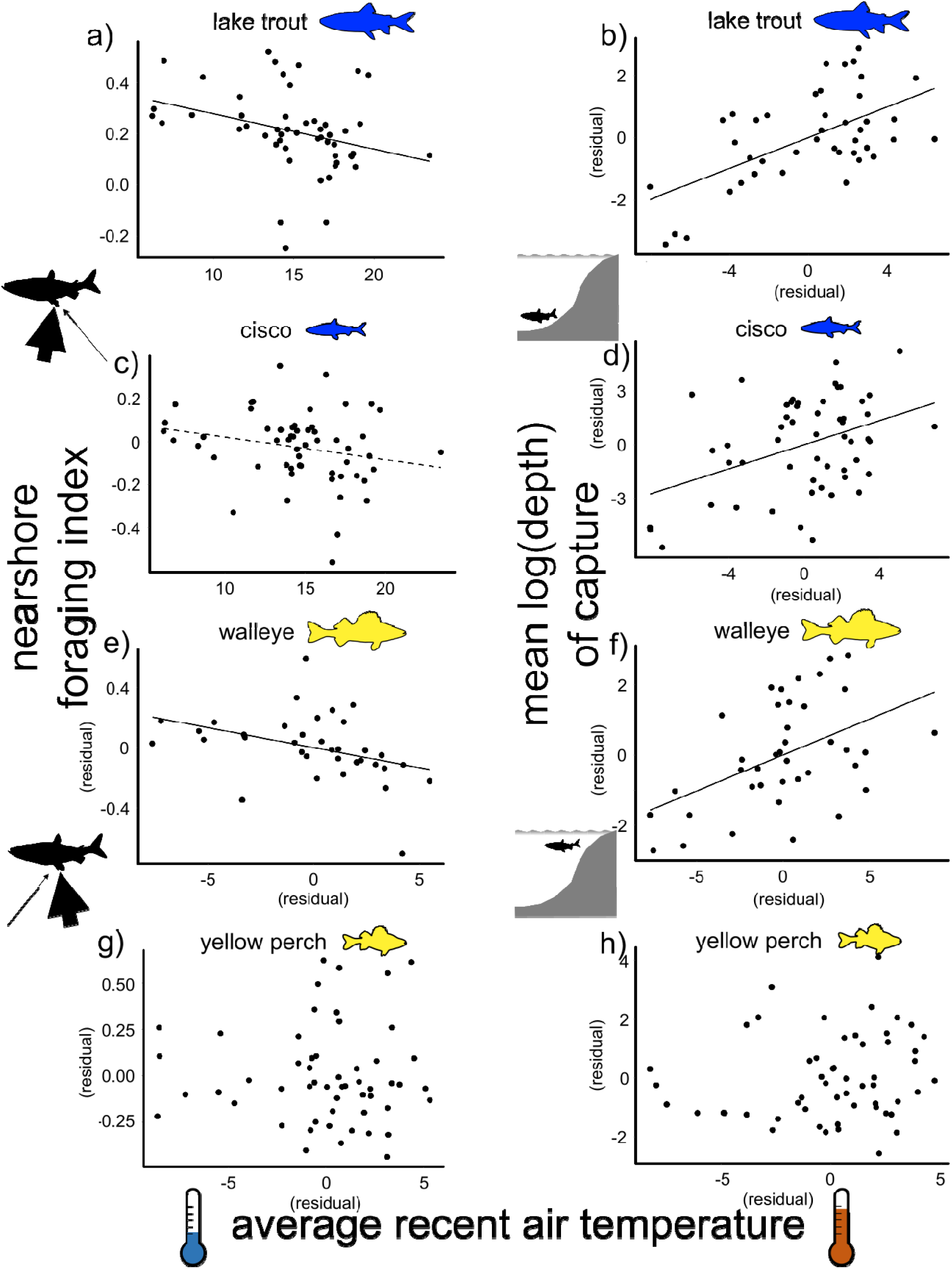
Nearshore foraging index and the mean log(depth) of capture of lake trout (a and b), cisco (c and d), walleye (e and f), and yellow perch (g and h) in lakes across a gradient of average recent air temperature. A solid line indicates a p-value less than 0.05, and a dashed line indicates a p-value less than 0.1. Regression model details can be found in Table S3. Fish silhouettes were taken from PhyloPic (http://www.phylopic.org).

## Results

### Food Web Metric Calculations using Stable Isotopes

We found strong evidence that lake food web structure varied across an approximately 7° C climate gradient. Three of the four species showed evidence of changes in nearshore feeding with increased temperature. Both top predators (lake trout and walleye) showed a significant decrease in the nearshore carbon index with increasing average recent air temperature, and the cold-adapted cisco showed a similar marginally significant decrease (Fig. 2a, c, and e, Table S3). The cool-adapted yellow perch showed no significant relationship between average recent air temperature and the nearshore carbon index (Fig. 2g, Table S3). Lake trout showed a significant increase in trophic position with increased average recent air temperature (Fig. 2b, Table S3), but cisco, walleye and yellow perch showed no significant relationship between trophic position with increasing average recent air temperature (Fig. 2d, Table S3).

**Figure 2.**
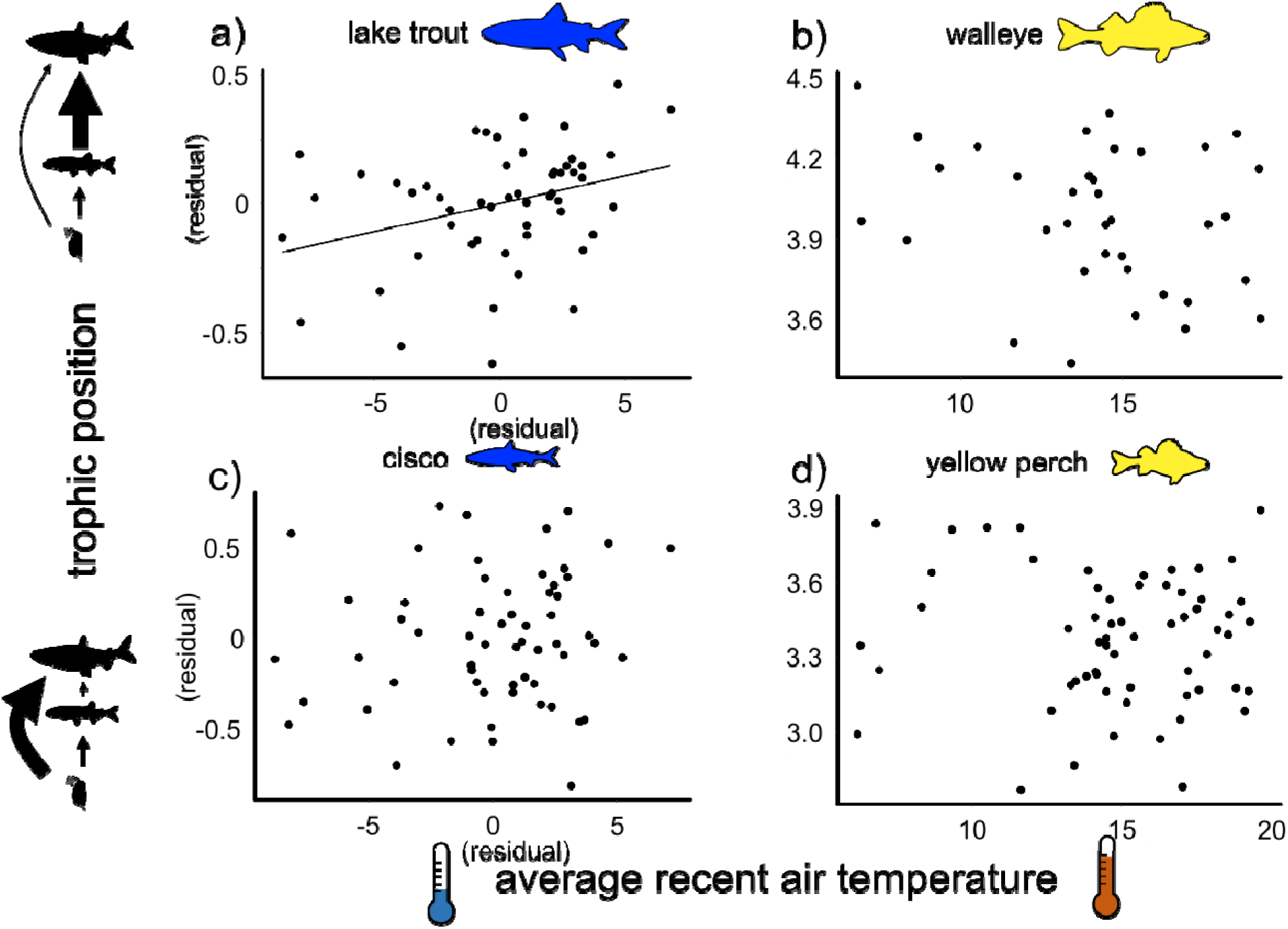
Trophic position of lake trout (a), walleye (b), cisco (c), and yellow perch (d) in lakes across a gradient of average recent air temperature. A solid line indicates a p-value less than 0.05, and a dashed line indicates a p-value less than 0.1. Regression model details can be found in Table S3. Fish silhouettes were taken from PhyloPic (http://www.phylopic.org).

### Behaviour Metrics and Biomass Index Using Catch-per-unit-effort

In agreement with changes in nearshore carbon index, the same three species (lake trout, walleye, and cisco) showed strong evidence of reduced nearshore habitat use. All three species showed a significant increase in mean depth of capture with increasing average recent air temperature (Fig. 3a, c, and e, Table S3). Consistent with these results, lake trout and cisco showed a significant decrease in probability of nearshore presence and walleye showed a marginally significant increase in probability of offshore presence with increasing average recent air temperature (see Supplementary Information). In contrast, yellow perch showed no relationship between either mean depth of capture or probability of offshore presence and average recent air temperature (Fig. 3g and inset, Table S3).

**Figure 3.**
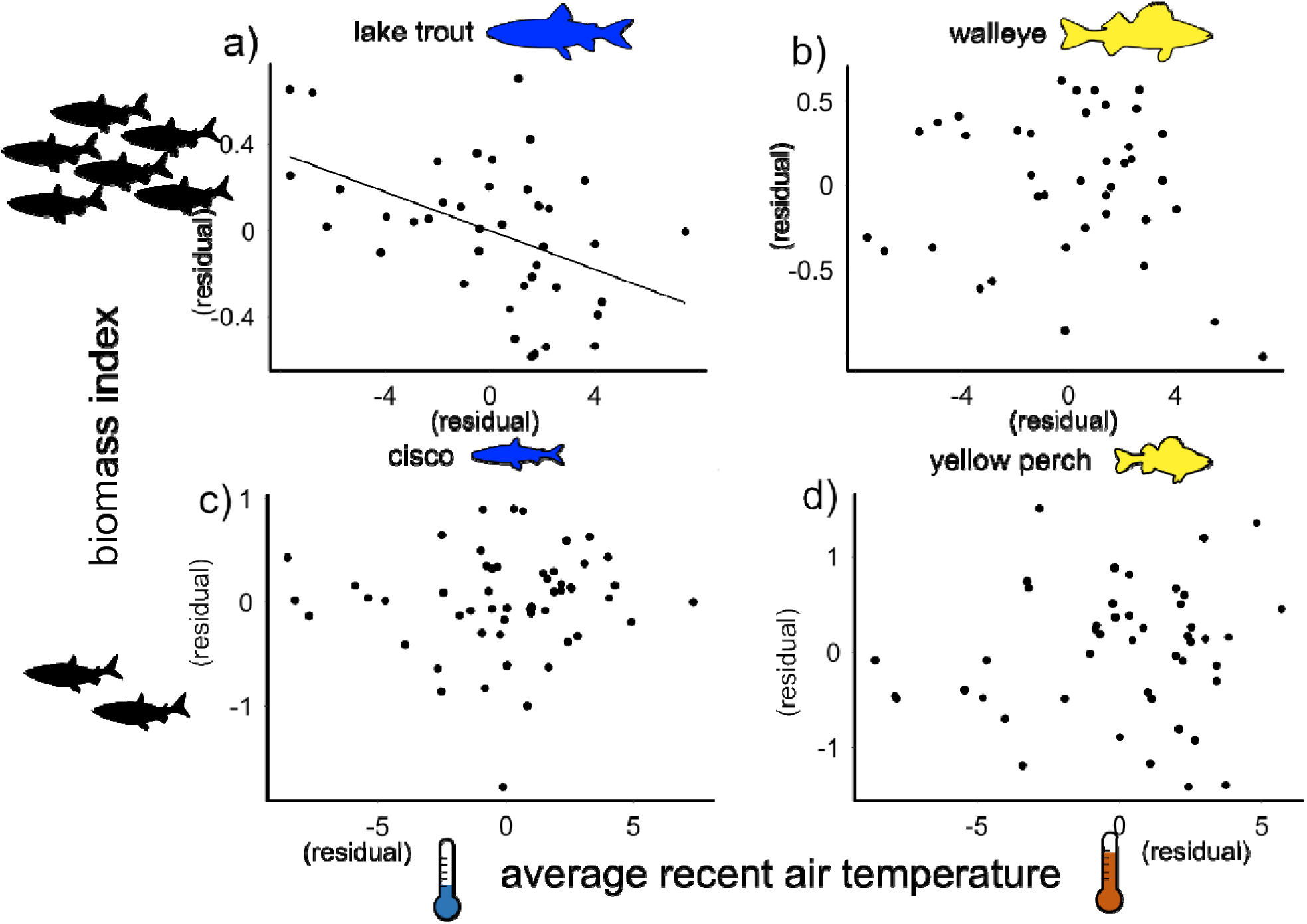
The log-transformed CUE biomass index for lake trout (a), walleye (b), cisco (c), and yellow perch (d) in lakes across a gradient of average recent air temperature. A solid line indicates a p-value less than 0.05, and a dashed line indicates a p-value less than 0.1. Regression model details can be found in Table S3. Fish silhouettes were taken from PhyloPic (http://www.phylopic.org).

**Figure 4:**
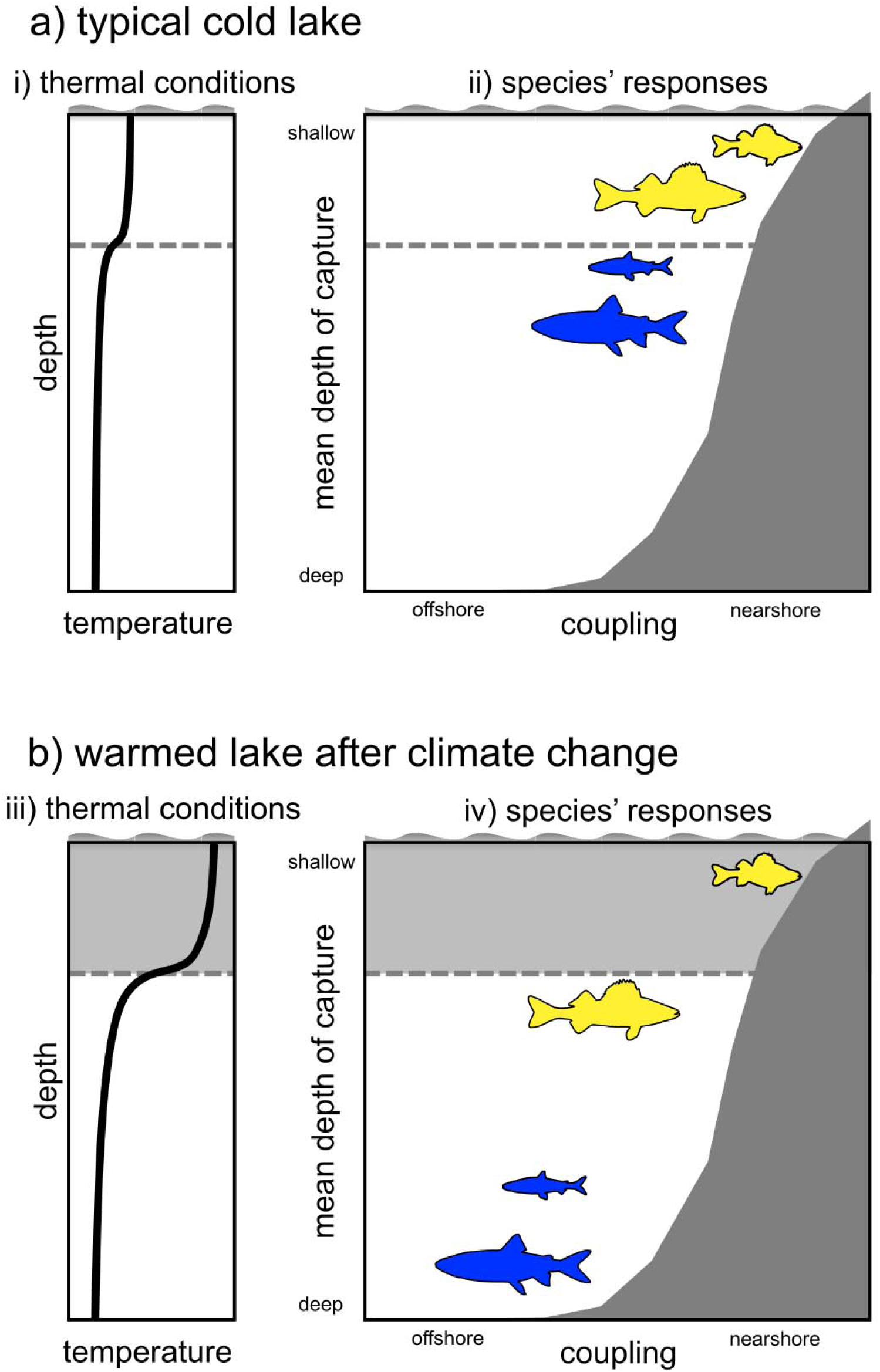
Schematic of the effects of warming on four widespread and abundant fish species of boreal shield lake food web: lake trout, cisco, walleye, and yellow perch. The x axis indicates shifts in the degree of nearshore foraging based on δ^13^C signatures. The y axis indicates shifts in habitat use in terms of the mean depth of capture. Lake trout, cisco, walleye all show shifts towards offshore foraging and offshore habitat use with increased temperature. Fish silhouettes were taken from PhyloPic (http://www.phylopic.org). Adapted from Tunney *et al.* 2014.

We found evidence of a reduction in biomass with increasing temperature for one species, lake trout, which showed a significant decrease in CUE biomass index with increasing average recent air temperature (Fig. 3b, Table S3). In contrast, cisco, walleye, and yellow perch showed no significant relationship between average recent air temperature CUE biomass index (Fig. 3d, f and h, Table S3).

## Discussion

Understanding how climate change is altering whole food webs is a challenging but critical endeavor for understanding the responses of whole ecosystems. Previous research showed that lake trout (*Salvelinus namaycush*), a generalist top predator in lake ecosystems, responds to increased temperature by reducing its nearshore habitat and resource use, and its degree of omnivory (Guzzo et al., 2017; T. D. Tunney et al., 2014), thus forming a flexible ‘generalist module’ (McMeans et al., 2016). Here, we extend that research to examine the impacts of climate warming on four key species that represent different key roles and trophic levels in boreal lake food webs: lake trout, walleye (*Sander vitreus*), cisco (*Coregonus artedi*), and yellow perch (*Perca flavescens*). Our stable-isotope-based food-web metrics and our catch-per-unit-effort-based behavioural metrics show a consistent response of species across multiple trophic levels to warming to nearshore-derived carbon and offshore habitat use. In fact, temperature was the most significant variable for predicting food web structure in this study (see Supplementary Information). By harnessing the variation in food web structure across natural temperature gradients, we have detected consistent food web responses to warming in boreal shield lake ecosystems.

We found strong evidence of foraging and behavioural responses to increased temperature in three of the four key food web members that we examined. As expected, our cold-adapted species, the predatory lake trout and the planktivorous cisco, showed decreased nearshore feeding and decreased near-shore habitat use with increasing temperatures. Our results are consistent with the thermally accessibility hypothesis that has been previously shown for lake trout (Guzzo et al., 2017; T. D. Tunney et al., 2014). Because offshore fish communities are largely comprised of species that prefer cold water (Coker et al., 2001; Hasnain et al., 2013), these thermal constraints likely operate on whole offshore fish communities, causing them to respond uniformly to warming. We found that the cool-adapted piscivorous walleye showed behavioural and foraging responses through decreased coupling and decreased nearshore habitat use with higher temperature, but in contrast the cool-adapted mesopredator yellow perch showed no such response. Since walleye and yellow perch have similar thermal tolerances (Hasnain *et al*. 2013), the reason for the different responses of these species is unclear. Despite the lack of response in yellow perch, the foraging responses of these fish species strongly indicate that boreal shield lake food webs will collectively flex towards offshore resource and habitat use in response to warming.

We found evidence that a reduction in nearshore coupling with warming may impact the biomass of lake trout. The biomass of lake trout decreased significantly with increased temperature, but none of the other species showed this pattern (Fig. 3). Such reductions in biomass with increasing temperature may be a consequence of a reduction in overall energy availability due to decreased nearshore coupling. Mobile top predators that garner energy from multiple spatially distinct macrohabitats are expected to have elevated biomass as a result of a higher overall resource availability, producing relatively top-heavy food webs (i.e., non-Eltonian, Mccauley *et al.* 2018; Woodson *et al.* 2018). Therefore, reduced access to the nearshore energy pathway in warm lakes could therefore reduce overall energy availability for lake trout and so lower lake trout biomass and population density (Guzzo et al., 2017; T. Tunney et al., 2018). A reduction in lake trout biomass across could subsequently reduce the top-down control of lake trout on cisco, their preferred prey. Reduced top-down control by lake trout in warmer lakes in conjunction with a bottom-up reduction in their biomass from reduced access to nearshore resources may result in no discernable trend in standing biomass of cisco with temperature, consistent with our results here. Changes in trophic biomass structure and concomitant changes in top-down control are one possible consequence of the flexes in lake food webs towards the offshore with warming. If climate change broadly decouples predators from multiple resource pools, changes in biomass structure may be a widespread signatures of climate change, possibly altering top-down effects within ecosystems (K. S. McCann, Rasmussen, & Umbanhowar, 2005; McCauley et al., 2018).

Our results demonstrate that many mobile generalist consumers at trophic levels and throughout compartments drive flexibility throughout food webs in boreal shield lakes. Our work builds on previous studies that have started to expose the role that single generalist top predators species play in determining food web responses to climate change in many ecosystems (Barton & Schmitz, 2009; Eloranta et al., 2016; Woodward, Perkins, & Brown, 2010). The presence of flexible foragers at various trophic levels is consistent with both recent ecological theory that predicts large, mobile generalists ought to rapidly respond to environmental change (Rooney & McCann, 2012), and is consistent with empirical evidence of flexible foraging from various ecosystems (Edmunds, Laberge, & McCann, 2016; Gonçalves, Azeiteiro, Pardal, & De Troch, 2012; Held, Mendl, Devereux, & Byrne, 2002; Rooney et al., 2008; Schmitz, Beckerman, & O’Brien, 1997; Shochat, Lerman, Katti, & Lewis, 2004; Taipale, Kankaala, Tiirola, & Jones, 2008; Vander Zanden & Vadeboncoeur, 2002). The presence of responsive foraging throughout diverse food webs rather than just at the top of food chains would make the generalist module the type of remarkably scale-invariant feature of food web architecture that has long interested ecologists (Sugihara, Schoenly, & Trombla, 1989).

The presence of responsive consumers across trophic levels and throughout food web compartments has strong implications for food web stability. Responsive foraging by generalists can be potently stabilizing in food webs (McMeans et al., 2016; D M Post, Conners, & Goldberg, 2000; Rip, McCann, Lynn, & Fawcett, 2010; Rooney, McCann, Gellner, & Moore, 2006). Generalist consumers can average variability in their resources (akin to the portfolio effect of Tilman *et al.* (1998)) by rapidly responding to resource variability in space and time by shifting away from low-density resources and towards high density resources, impeding that variability from emanating throughout food webs (Valdovinos, Ramos-Jiliberto, Garay-Narváez, Urbani, & Dunne, 2010). This switching ability also increases trophic redundancy, which can prevent secondary extinctions (Borrvall, Ebenman, & Jonsson, 2000; Sanders, Thébault, Kehoe, & Frank van Veen, 2018). If a broad suite of organisms can make rapid, smart foraging responses to environmental change, they may accentuate the stabilizing effect of a single generalist module. The presence of many responsive consumers increases stability in complex food web models (Kondoh, 2010; Valdovinos et al., 2010). This reveals a potential powerful, repeated stabilizing mechanism throughout food webs that Levin (1998) argues is fundamental to persistence of ecosystems. However, when generalist consumers decouple macrohabitats to avoid entering physiologically taxing habitats, their ability to respond to consumer densities is limited, undermining their stabilizing potential (Guzzo et al., 2017; Schmitz & Barton, 2014; T. D. Tunney et al., 2014). Thus, the decoupling of habitats with warming may be widespread across ecosystems, forcing the food webs of many ecosystems to flex with climate change in ways that alter ecosystem stability worldwide.

Our work here adds a key piece to the mounting evidence that generalist consumers likely to play a vital role in determining how climate change rewires food webs (Bartley et al., n.d.). Generalist consumers positioned throughout food webs may ultimately reorganize whole ecosystems in response to climate change (Blanchard, 2015; McMeans et al., 2016). Determining both the prevalence and positions of these responsive generalist consumers and the factors that constrain these species responses are critical tasks for predicting the myriad ways that food webs will adapt with a changing climate. Importantly, our results suggest that these species throughout food webs contribute to adaptive capacity—the ability of an ecosystem to change in response to changing conditions—a key part of the resilience of ecosystems in the face of environmental variation (McMeans et al., 2016). Although we have showed that lake food webs flex at many axes, it is unclear when this adaptive capacity will be exceeded and entire ecosystems collapse. This uncertainty highlights the clear need to forecast major changes in food web structure. Monitoring the behaviour and foraging of responsive consumers is a promising way to track changes in food web structure and give prescient warnings of future drastic changes in ecosystem dynamics (Bartley et al., n.d.; Velarde, Ezcurra, & Anderson, 2013). This makes responsive generalist consumers critical to understanding of how ecosystems will fare through climate change.

## Acknowledgements

This work was supported by the National Science and Engineering Research Council of Canada through a Discovery Grant to K.S.M., a Strategic Grant to K.S.M., B.J.S. and N.P.L., and a Canada Graduate Scholarship to T.J.B. We thank K. Armstrong, J. Wright and many other biologists and technical staff at the Ontario Ministry of Natural Resources and Forestry and the Broad-scale Fisheries Monitoring Program. We also thank M. Berger, I. Farahbakhsh, B. Graham, B. Heavenrich for assistance with gathering and organizing climate data. D. Schindler, M. Granados, M. Guzzo, and B. Robinson provided helpful comments on an early draft of this manuscript.

## Author Contributions

T.J.B. and K.S.M. conceived and designed this study. T.J.B. and T.D.T. collected field data and conducted lab work. T.J.B. analyzed data and wrote the initial draft of the manuscript. T.J.B., T.D.T., N.P.L., B.J.S., R.H.H., and K.S.M. all contributed substantially to writing and revisions.

## Data Accessibility

Upon acceptance, our data will be archived on Zenodo, https://zenodo.org/

