## Supplementary Information for "Multiple species drive flexible lake food webs with warming"

Timothy J. Bartley^1,2^*, Tyler D. Tunney^3,4^, Nigel P. Lester^5,6^, Brian J. Shuter^5,6^, Robert H. Hanner^2,7^, and Kevin S. McCann^2^

^1^Department of Biology, University of Toronto Mississauga, 3359 Mississauga Road, Mississauga, Ontario, Canada, L5L 1C6

^2^Department of Integrative Biology, University of Guelph, 50 Stone Road East, Guelph, Ontario, Canada N1G 2W1

^3^Gulf Fisheries Centre, Fisheries and Oceans Canada, Moncton, New Brunswick E1C 9B6, Canada

^4^Center for Limnology, University of Wisconsin–Madison, Madison, WI, USA 53706

^5^Science and Research Branch, Ontario Ministry of Natural Resources, Peterborough, Ontario, Canada K9J 7B8

^6^Department of Ecology and Evolution, University of Toronto, Toronto, Ontario, Canada M5S 3G5

^7^Biodiversity Institute of Ontario, University of Guelph, 50 Stone Road East, Guelph, Ontario, Canada N1G 2W1

*correspondence to T.J.B.

**Supplementary Methods**

*Estimates of Baseline Stable Isotopic Signatures*

For the two lakes that were missing nearshore baseline isotopic signatures and for the five lakes that were missing offshore baseline isotopic signatures, we estimated the missing isotopic signatures of using the available baseline and simple linear regression between the average baseline signatures across lakes. The equations for the linear regression models used to estimate missing nearshore baselines (i.e., snails and aquatic insect larvae) and offshore baselines (i.e., mussels and zooplankton) were

$$\delta^{13}C_{nsb est}=-6.91152+0.503579 \times\delta^{13}C_{osb meas}$$

$$\delta^{13}C_{osb est}=-25.0199+0.181308 \times\delta^{13}C_{nsb meas}$$

$$\delta^{15}N_{nsb est}=0.089087+0.785173 \times\delta^{15}N_{osb meas}$$

$$\delta^{15}N_{osb est}=1.234742+0.722821\times\delta^{15}N_{nsb meas}$$

where $\delta^{13}C_{nsb est}$is the estimated nearshore δ^13^C signature, $\delta^{13}C_{osb meas}$ is the measured average δ^13^C signature, $\delta^{13}C_{osb est}$is the estimated offshore δ^13^C signature, $\delta^{13}C_{nsb meas}$ is the measured average nearshore δ^13^C signature, $\delta^{15}N_{nsb est}$is the estimated nearshore δ^15^N signature, $\delta^{15}N_{osb meas}$ is the measured average δ^15^N signature, $\delta^{15}N_{osb est}$is the estimated offshore δ^15^N signature, $\delta^{15}N_{nsb meas}$ is the measured average nearshore δ^15^N signature. The R^2^ for the δ^13^C regression models was 0.09254, and the R^2^ for the δ15N regression models was 0.5675.

*Logistic Regressions for Nearshore or Offshore Presence*

We supplemented our analyses of mean depth of capture using the probability of nearshore or offshore capture each species because the stratified sampling used by Sandstrom et al. (2013) may not adequately capture changes in habitat use by these species. We considered each species to be present in the nearshore (6m of water or shallower) or offshore (greater than 6m of water) if the CUE of that species in that habitat was > 0. We calculated the proportion nearshore CUE for each species in each lake as the sum of the CUE for that species in depth strata < 6 m in that lake divided by the total CUE of that species for that lake. Similarly, we calculated the proportion offshore CUE for each species in each lake as the sum of the CUE for that species in depth strata in water > 6 m in that lake divided by the total CUE of that species for that lake. Because offshore and nearshore species were always caught in their associated habitats but not always caught in their non-preferred habitat, we constructed logistic regression models of the nearshore catch probability to examine offshore species’ behaviour and the offshore catch probability to examine nearshore species’ behaviour. Logistic regression models were weighted with the proportion of each species’ catch in their non-preferred habitat + 1.

**Supplementary Results**

With increasing growing degree days, lake trout showed a significant decrease in proportion of nearshore carbon, a marginally significant increase in trophic position, no significant relationship with mean depth of capture, a significant decrease in nearshore catch probability, and a significant decrease in CUE biomass index. With increasing growing degree days, cisco showed a significant decrease in proportion littoral carbon, a significant increase in trophic position, no significant relationships with nearshore catch probability, mean depth of capture, or CUE biomass index. With increasing growing degree days, walleye showed a marginally significant decrease in proportion littoral carbon, no significant relationship with trophic position, mean depth of capture, or offshore catch probability, and a significant decrease in CUE biomass index. With increasing growing degree days, yellow perch showed no significant relationship with proportion littoral carbon or trophic position, a significant increase in mean depth of capture, no relationship with offshore catch probability, and a marginally significant increase in CUE biomass index.

**Supplementary Tables**

**Table S1.** Summary of the 7 temperature, physical and chemical lake attributes for lakes used in our study.

| Variable | number of lakes | min | max | mean | sd |
| --- | --- | --- | --- | --- | --- |
| Average Recent Air Temperature in °C | 66 | 6.18 | 23.5 | 14.9 | 3.49 |
| Growing Degree Days > 5 °C | 66 | 1436 | 2214 | 1645 | 147 |
| Surface Area in ha | 66 | 48.0 | 11620 | 2680 | 2737 |
| Shoreline Development Index | 59 | 1.70 | 14.95 | 6.10 | 3.47 |
| Mean Lake Depth in m | 66 | 3.80 | 39.0 | 12.9 | 6.89 |
| Secchi in Summer in m | 63 | 0.80 | 8.60 | 4.08 | 1.69 |
| Total Phosphorous in μg•L^-1^ | 59 | 3.20 | 20.8 | 8.37 | 3.94 |

**Table S2.** The thermal guild, trophic level, average CUE in lakes where present, and the number of lakes in which each species in present for the four boreal shield fish species used in this study and for all other species combined. Data are for 59 lakes sampled by the Ontario Ministry of Natural Resources and Forestry’s Broad-scale Fisheries Monitoring (BSM) Program (Sandstrom *et al.* 2013). Thermal guild and trophic level are taken from Coker *et al.* 2001 and Hasnain *et al.* 2013.

| common name | scientific name | thermal guild | trophic level | average CUE | number of lakes |
| --- | --- | --- | --- | --- | --- |
| lake trout | *Salvelinus namaycush* | cold | predator | 9.62 | 47 |
| cisco | *Coregonus artedi* | cold | intermediate consumer | 32.30 | 58 |
| walleye | *Sander vitreus* | cool | predator | 25.55 | 41 |
| yellow perch | *Perca flavescens* | cool | intermediate consumer | 92.31 | 57 |
| all other species |  | various | various | 217.12 | 59 |

**Table S3.** Regression model (linear or logistic) summaries for nearshore carbon index, trophic position, catch probability in the non-preferred habitat, mean depth of capture, and the CUE biomass index (catch-per-unit-effort in kg) of lake trout, cisco, walleye, and yellow perch with average recent air temperature in °C (T).

| Species | Dependent Variable | Non-Temperature Variable | Model | adj. R^2^ | df | temperature coefficient p-value | overall p-value |
| --- | --- | --- | --- | --- | --- | --- | --- |
| lake trout | nearshore carbon index | none | *y* = 0.416021 – 0.013817(T) | 0.08346 | 1, 49 | 0.0225 | *NA* |
|  | trophic position | log_10_(mean lake depth in m) (LMD) | *y* = 3.898934 + 0.021658(T) + 0.103122(LMD) | 0.1516 | 2, 50 | 0.00764 | 0.006161 |
|  | mean depth of capture | log_10_(lake surface area in ha) (LSA)  log_10_(mean lake depth in m) (LMD) | *y* = 3.48763 + 0.24655(T) + 1.70181(LSA) + 0.79042(LMD) | 0.573 | 3, 38 | <0.0001 | <0.0001 |
|  | nearshore presence | log_10_(lake surface area in ha) (LSA) | log($\frac{y}{1-y})$ = 10.1488 – 0.2524(T) + 2.2319(LSA) | *NA* | 2, 43 | 0.01892 | 0.007497 |
|  | CUE biomass index | log_10_(lake surface area in ha) (LSA)  log(Secchi depth in m) (LSD) | *y* = 2.74427 – 0.03245(T) – 0.28596(LSA) | 0.2655 | 2, 40 | 0.01508 | 0.0007872 |
| cisco | nearshore carbon index | none | *y* = 0.125846 – 0.010469(T) | 0.03375 | 1, 54 | 0.09317 | *NA* |
|  | trophic position | log_10_(mean lake depth in m) (LMD) | *y* = 2.64522 + 0.01299(T) + 0.24621 (LMD) | 0.09171 | 2, 56 | 0.3599 | 0.02533 |
|  | mean depth of capture | log(Secchi depth in m) (LSD)  log(total phosphorous in μ·L^−1^) (LTP) | *y* = -8.6812 + 0.5749(T) + 5.2551(LMD) + 3.9327(LTP) | 0.3351 | 3, 51 | 0.003810 | <0.0001 |
|  | nearshore presence | log_10_(lake surface area in ha) (LSA)  log(shoreline development index) (LSDI)  log_10_(mean lake depth in m) (LMD) | log($\frac{y}{1-y})$ = 11.3375 – 0.6778(T) –1.5685(LSA) – 0.6899(LSDI) – 3.2943(LMD) | *NA* | 4, 56 | 0.00222 | <0.0001 |
|  | CUE biomass index | log(shoreline development index) (LSA)  log(Secchi depth in m) (LSD)  log(total phosphorous in μ·L^−1^) (LTP) | *y* = – 0.452757+ 0.009392(T) + 0.228017 (LSA) + 0.290588(LSD) + 0.368038(LTP) | 0.08951 | 4, 49 | 0.6640 | 0.07168 |
| walleye | nearshore carbon index | log_10_(mean lake depth in m) (LMD) | *y* = 0.91052 – 0.02655(T) – 0.09965(LMD) | 0.1938 | 2, 33 | 0.0201 | 0.01083 |
|  | trophic position | none | *y* = 4.27798 + 0.02102(T) | 0.04287 | 1, 34 | 0.118 | *NA* |
|  | mean depth of capture | log_10_(lake surface area in ha) (LSA) | *y* = – 1.55047 + 0.20567(T) + 1.18090(LSA) | 0.2943 | 2, 36 | 0.0023 | 0.0007119 |
|  | offshore presence | none | log($\frac{y}{1-y})$ = –10.5347 + 0.4487(T) + 2.4169(LSA) | *NA* | 2, 38 | 0.07190 | 0.06644 |
|  | CUE biomass index | log_10_(mean lake depth in m) (LMD) | *y* = 3.4664033 0.0007749(T) – 1.9555856(LMD) | 0.3882 | 2, 36 | 0.977 | <0.0001 |
| yellow perch | nearshore carbon index | log_10_(lake surface area in ha) (LSA)  log(total phosphorous in μg·L^−1^) (LTP) | *y* = 0.3653330 + 0.0110075(T) + 0.0767033(LSA) + 0.0866055(LTP) | 0.08208 | 3, 50 | 0.9891 | 0.06387 |
|  | trophic position | none | *y* = 3.441276 – 0.004554(T) | -0.0134 | 1, 59 | 0.65 | 0.6502 |
|  | mean depth of capture | log(Secchi depth in m) (SD) | *y* = 1.96064 + 0.07213(T) + 0.96622(SD) | 0.1272 | 2, 51 | 0.2535 | 0.01168 |
|  | offshore presence | log_10_(lake surface area in ha) (LSA)  log(Secchi depth in m) (LSD) | log($\frac{y}{1-y})$ = 1.58785 + 0.03013(T) | *NA* | 1, 55 | 0.7740 | *NA* |
|  | CUE biomass index | log_10_(mean lake depth in m) (LMD) | *y* = 0.9181 + 0.0114(T) + 0.6075(LMD) | 0.008427 | 2, 52 | 0.701 | 0.3008 |
